# A Tetravalent Dengue Virus-like Particle Vaccine Induces High levels of Neutralizing Antibodies and Inhibits Dengue Replication in Non-Human Primates

**DOI:** 10.1101/2023.10.16.562563

**Authors:** Daniel Thoresen, Kenta Matsuda, Akane Urakami, Mya Myat Ngwe Tun, Takushi Nomura, Meng Ling Moi, Yuri Watanabe, Momoko Ishikawa, Trang Thi Thu Hau, Hiroyuki Yamamoto, Yuriko Suzaki, Yasushi Ami, Jonathan F Smith, Tetsuro Matano, Kouichi Morita, Wataru Akahata

## Abstract

Dengue virus (DENV) represents a significant global health burden, with 50% of the world’s population at risk of infection, and there is an urgent need for next-generation vaccines. Virus-like particle (VLP)-based vaccines, which mimic the antigenic structure of the authentic virus but lack the viral genome, are an attractive approach. Here we describe a dengue VLP (DENVLP) vaccine which generates a robust and long-lasting neutralizing antibody response against all four DENV serotypes in non-human primates. Importantly, DENVLP vaccination produced no ADE response against any of four DENV serotypes. Finally, we demonstrate in a non-human primate challenge model that DENVLP vaccination substantially reduces viral replication. We also transfer the purified IgG from the immunized monkeys into immunodeficient mice, where they protect against subsequent lethal dengue virus challenge, indicating a humoral mechanism of protection. These results indicate that a DENVLP vaccine is a safe and effective vaccine candidate.

**One Sentence Summary:** Immunization of non-human primates with a tetravalent dengue VLP vaccine induces high levels of neutralizing antibodies and reduces the severity of infection for all four dengue serotypes.

## Introduction

Dengue is a viral disease that infects nearly 400 million people worldwide each year, and in severe cases causes dengue hemorrhagic fever (DHF), which is responsible for approximately 10,000 deaths each year^1,2^. Dengue virus (DENV) is spread by both *Aedes aegypti* and *Aedes albopictus* mosquitos, which are endemic to approximately 100 countries across all 6 habitable continents^3,4^. Furthermore, in the coming decades the range of both vector hosts will likely increase due to climate change, further increasing the population at risk of infection^5^. No therapeutic drug to treat dengue infection is currently licensed, which makes an effective vaccine essential to meeting this global health threat.

A significant obstacle to developing a dengue vaccine is the phenomenon known as antibody-dependent enhancement (ADE)^6^. There are four closely related but antigenically distinct DENV serotypes (DENV1-4). An individual’s first DENV infection is typically mild or asymptomatic. However, a subsequent infection with a different DENV serotype may result in more severe disease due to ADE. ADE occurs when pre-existing anti-DENV antibodies against one DENV serotype form a DENV-antibody immunocomplex which enhances cellular entry into Fc-receptor-bearing cells and worsens the infection rather than preventing it^7^. Importantly, ADE can also be caused by the antibodies generated by a vaccine, causing a person’s first dengue exposure to lead to more severe illness. Therefore, to prevent ADE risk, any dengue vaccine must produce strong neutralizing antibody (NAb) responses against all four serotypes simultaneously.

To date, two dengue vaccines have been licensed and approved. The first, Dengvaxia, is a live-attenuated tetravalent vaccine developed by Sanofi Pasteur and approved in 2016. However, long-term safety and efficacy studies revealed variable efficacy based on age and dengue sero-status at the time of vaccination, as children 2-9 years old at initial vaccination exhibited an elevated risk of severe dengue infection and hospitalization relative to children in the control group^8,9^. Due to this risk, Dengvaxia is only recommended for persons 9 to 16 years of age in the areas of high dengue-endemicity (>70% seroprevalence)^10,11^. The second approved vaccine, Qdenga (TAK-003), is a live-attenuated tetravalent vaccine developed by Takeda, and has been recommended for use in children age 6-16 in areas of high dengue disease burden^12^. The phase 3 clinical trial results indicated that vaccine recipients who were seronegative prior to immunization were more likely to develop DENV3 or DENV4 infections, and more likely to be hospitalized with DENV3 than seronegative individuals receiving placebo^13^. Although the precise mechanism behind the elevated risk in seronegative recipients is unknown, one possibility is that it is due to ADE resulting from insufficient antibody production against one or more serotypes. With a live-attenuated vaccine, the replication of some attenuated DENV sterotype may be inhibited by the replication of the other serotypes, leading to a lack of sufficient antigen production. An approved vaccine for seronegative individuals remains a major global health objective.

Virus-like particles (VLPs) are self-assembling structures composed of viral structural proteins without genomic DNA or RNA. VLP vaccines present a repetitive, high-density antigen profile that closely mimics the morphology of an authentic virus, making them highly immunogenic^14^. However, the absence of genomic information required to replicate makes them relatively safe even in young children and immunocompromised individuals. The ability to immunize young children has critical importance for reducing dengue morbidity as a study of DHF prevalence in Thailand demonstrated that infants <1 year old represent a significant proportion of severe DHF cases^15^. One hypothesis for the cause of severe dengue infection in infants is that it stems from ADE caused by maternal DENV antibodies acquired *in utero*. As the passively transferred maternal antibody titer wanes, the once protective antibody can result in enhanced dengue infection in the infant^16,17^. The live attenuated vaccines are not considered for immunization of infants due to safety issues and concerns that the presence of the maternal antibodies may negatively affect the replication of one or more attenuated serotypes, leading to an imbalanced response^18,19^. Immunocompromised individuals are another group at high risk of severe dengue infection, however live-attenuated vaccines are not recommended due to the risk of increased replication or genetic reversion^20,21^. VLP vaccines, on the other hand, are safe for immunocompromised persons because they lack the genetic material required for replication and reversion. These characteristics make a VLP vaccine attractive for protecting both high-risk groups.

In this study, we describe a tetravalent DENVLP vaccine and demonstrate that it induces a robust neutralizing antibody response against all four DENV serotypes for up to 1 year in non-human primates. Furthermore, ADE activity was not detected against any serotype throughout the 1-year timeframe. We also show that the anti-DENV neutralizing antibodies induced by this vaccine are capable of significantly reducing viral replication in rodents and non-human primates.

## Results

### A tetravalent DENV VLP vaccine generates robust and long-lasting immunity

We have previously described the generation of an envelope-modified tetravalent DENVLP vaccine^22^. The prM-E region of DENV1-4 were selected for VLPs due to its ability to spontaneously form the VLP structure as well as the ability to generate strong neutralizing antibody levels. In all four constructs, a mutation in the fusion loop (F108A) of envelope domain II (EDII) enabled a significant increase in VLP production without disrupting the vaccine’s ability to induce neutralizing antibodies (**Fig. 1A**). In the DENVLP2-4 constructs, the envelope domain III, stem, and transmembrane anchor (EDIII/ST/TM) of the native serotype were replaced with the EDIII/ST/TM of DENV1 (shown in blue) in order to increase vaccine production. This substitution was confirmed to not affect the ability of DENVLP2-4 to produce serotype-specific neutralizing antibodies in mice^22^.

**Figure 1.**
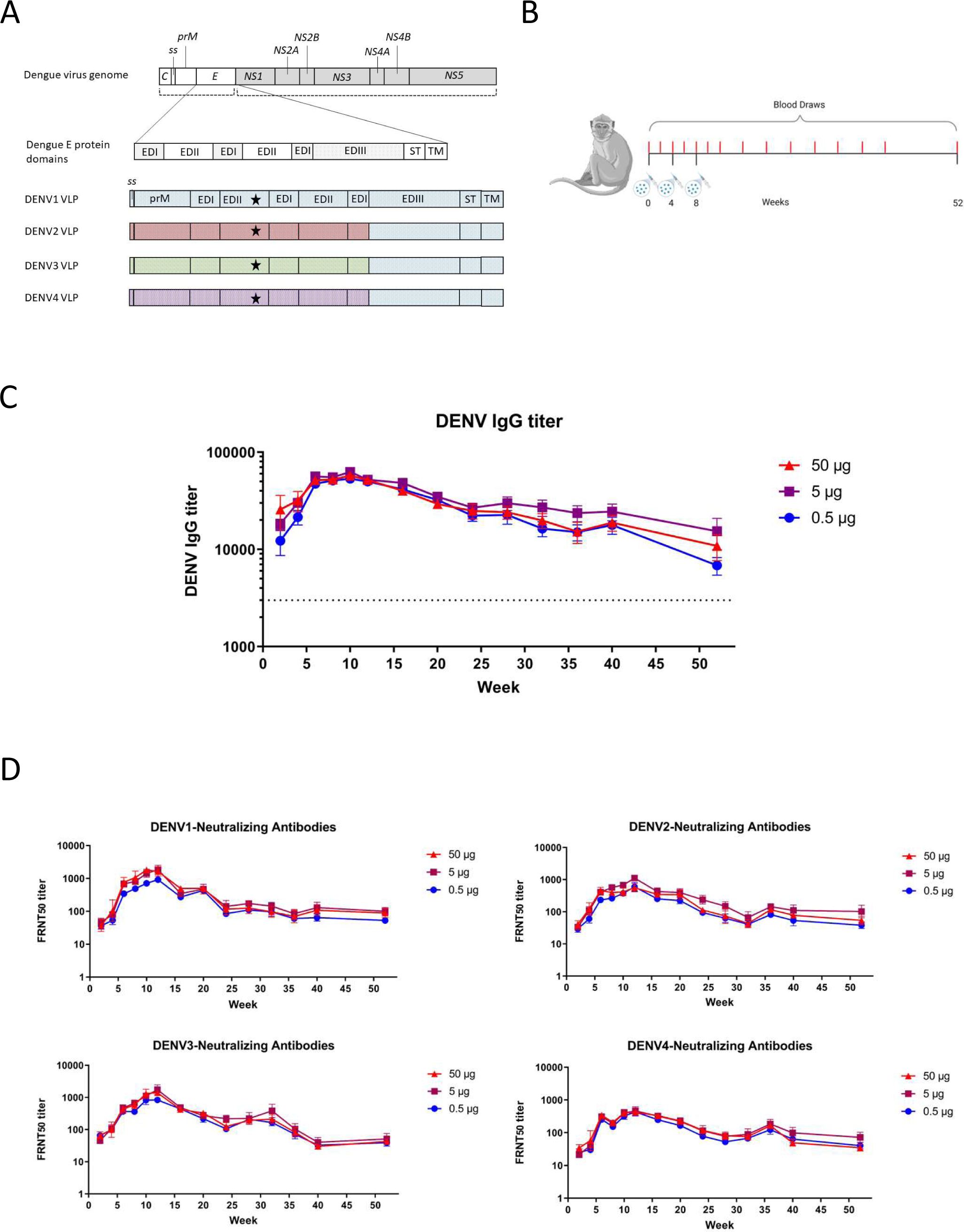
A Tetravalent DENVLP vaccine generates robust antibody response. **(A)** Schematic illustration of the DENV genome and DENV1-4 VLP expression vectors. C, capsid; ss, signal sequence; prM, precursor membrane; E, envelope; NS1-5, Non-structural proteins 1-5; EDI-III, envelope domains I-III; ST, stem; TM, transmembrane domain. Star symbol indicates location of F108A substitution in VLP expression constructs. **(B)** Graphic of macaque immunization experiment. Tetravalent DENVLP (0.5, 5 and 50 µg, n = 6 per dose group) adjuvanted with alum was administered at weeks 0, 4 and 8. Serum samples (shown in red) were taken every two weeks from weeks 0-12 and every four weeks from weeks 12-52. **(C)** The mean anti-DENV IgG titer for all three dose groups was determined by ELISA for each time point. Serum samples from each macaque and timepoint were diluted 1:1000, and after labeling with an HRP-conjugated secondary antibody and developed with substrate, the OD_492_ value for each sample was determined from a standard curve. The minimum threshold for a positive result, shown as the dotted line, is set to an IgG titer of 3000. **(D)** The mean neutralizing antibody titer for each dose group and time point as determined by FRNT_50_ assay. Serum samples from each macaque and timepoint were serially diluted and incubated with DENV serotypes 1-4 prior to inoculation of Vero cells. The endpoint dilution capable of neutralizing half of the viral foci relative to Vero cells incubated with DENV1-4 alone was recorded for each macaque and timepoint was recorded, and the mean value for each dose group and sample timepoint was plotted.

To better approximate the human immune response to DENVLP vaccination, we conducted immunogenicity and protection studies in non-human primates (NHPs), which have been previously shown to develop immune responses that closely resemble those seen in humans^17,18^. Cynomolgus macaques (*Macaca fascicularis)* were immunized with one of three doses (0.5, 5, or 50 µg total DENVLP, containing equal amounts of each serotype, n = 6 per group) of tetravalent DENVLP vaccine at 0, 4 and 8 weeks (**Fig. 1B**). The total IgG antibody titer against all four DENV serotypes was measured via ELISA (**Fig. 1C**), and the mean anti-DENV IgG titer at the peak (week 10) for the lowest vaccine dose (0.5 µg) was 53350 ± 4977, which was not significantly different from the mean IgG titer of 57510 ± 14314 for the highest dose (50 µg). By 52 weeks after the first immunization, the total anti-DENV IgG titer had declined to between 1/4^th^ and 1/8^th^ of the peak IgG titer observed depending on the dose of vaccine delivered, but the IgG titer for the highest and lowest doses of vaccine were still not statistically significant. The IgG titer for all three vaccine doses remained above the threshold of detection at the 52-week point, indicating that this vaccine regimen produced a humoral response up to 1 year and beyond.

To determine if the antibodies produced by the DENVLP vaccine could sufficiently neutralize all four dengue serotypes, we measured the neutralizing antibody (NAb) titer against each specific virus serotype (DENV1-4) in the serum of the same macaques. We observed substantial NAb titers generated to each DENV serotype, with peak neutralizing antibody titers observed at either week 10 or 12 (1848, 1120, 1732, and 462 against DENV1-4, respectively) for all vaccine doses against all four serotypes (**Fig. 1D**). By week 52, the NAb titer had waned from the peak (100, 102, 51, and 72 against DENV1-4, respectively), but remained above the threshold of detection, indicating that a neutralizing antibody response was maintained for up to 1 year or beyond following immunization. At most timepoints the difference in NAb titer between the highest and lowest dose were not statistically significant, suggesting that all three dose levels could produce a robust neutralizing response. Comparing the NAb titer between the four strains for all 18 immunized animals, the greatest fold difference in NAb titer observed was a 5-fold difference between DENV1 and DENV4 at week 8, however following the third vaccine dose the serotypes with the highest and lowest NAb titers varied from sample to sample (**Fig. S1**). This variation suggests that the vaccine does not produce an imbalanced neutralizing response that favors neutralization of one DENV serotype over the other three.

### Tetravalent DENVLP vaccination does not generate ADE for DENV serotypes 1-4

Immunization with dengue vaccines can potentially cause ADE by inducing both neutralizing and non-neutralizing antibodies. When the level of neutralizing antibodies is insufficient to fully neutralize virions, infection can be enhanced rather than suppressed. To examine whether the antibodies generated by DENVLP immunization of cynomolgus monkeys were capable of increasing infectivity via ADE, DENV1-4 virions were incubated with the indicated dilutions of either purified monoclonal neutralizing antibodies (Control) or serum from tetravalent DENVLP-immunized macaques, and then Fc-gamma receptor-expressing baby hamster kidney cells (BHK-FcγR) were infected with the antibody-incubated virions. With two purified monoclonal anti-dengue neutralizing antibodies, high levels of antibody suppressed the infection of BHK cells, resulting in a ratio of infectivity <1 when compared to virus-infected cells alone (**Fig. 2A**). However, as the monoclonal antibodies are diluted further, the ratio of infectivity rises above 1, demonstrating that dengue infection of the BHK-FcγR cells will be enhanced as antibody titer decreases. For DENVLP-immunized cynomolgus monkeys (n=18), a 1:10 serum dilution failed to generate ADE through 52 weeks (**Fig. 2B-E**), with levels consistently below the acceptable threshold for ADE. Since ADE did not occur at the low dilution (1:10) of sera fromany of the samples at any time point (in which absolute titers varied significantly), it is unlikely that immune enhancement would occur under *in vivo* conditions (e.g., with undiluted serum) up to one year after vaccination.

**Figure 2.**
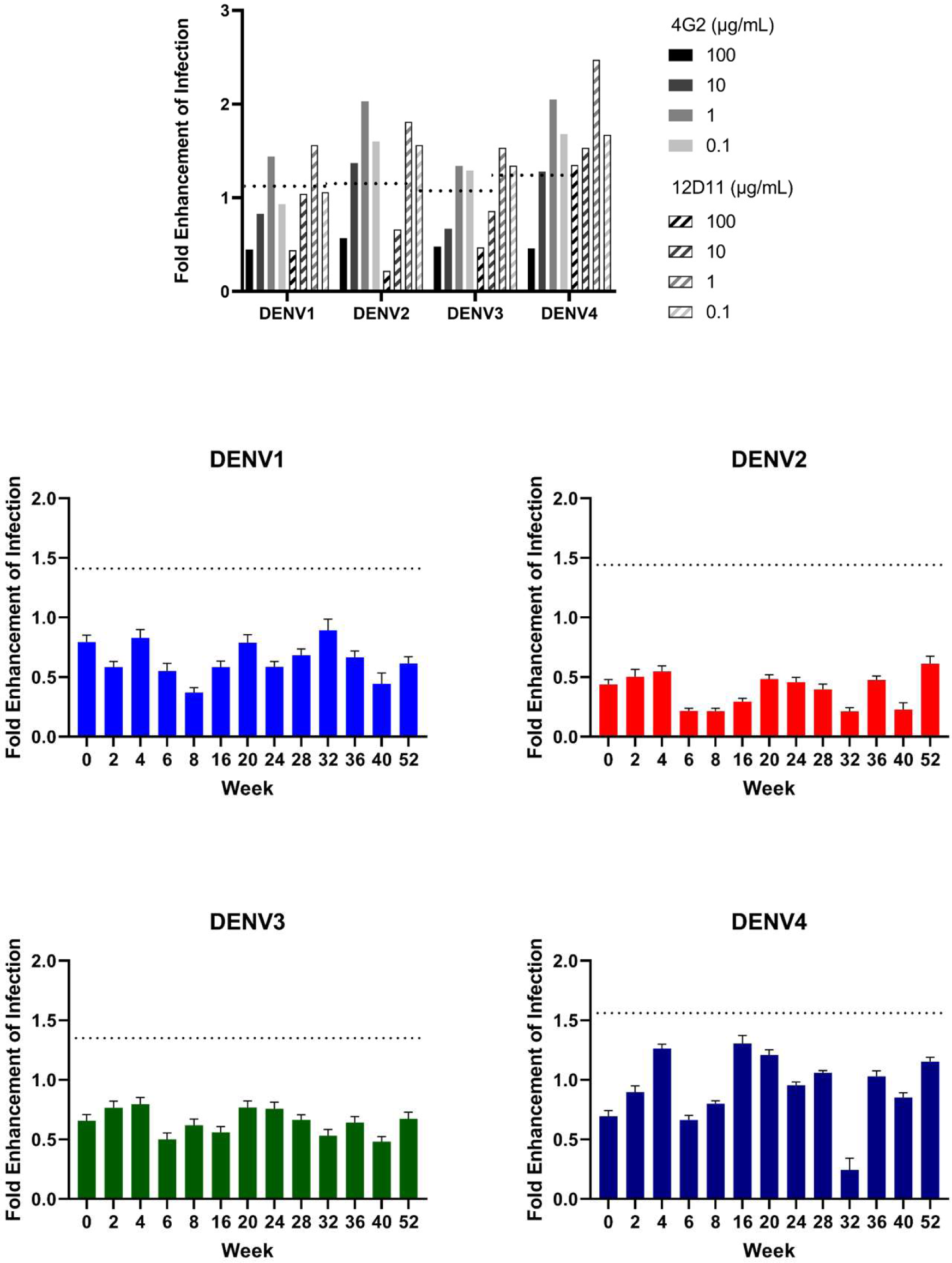
Dilution of neutralizing antibodies can cause ADE activity, but not for macaque sera diluted 1:10. **(A-E)** DENV1-4 were incubated with monoclonal antibodies at indicated dilutions **(A)** or macaque sera diluted 1:10 **(B-E)** prior to inoculation of FCγR-BHK cells. After two days, infected cells were counted, and the ratio of infected cells in serum- or antibody-incubated wells to infected cells in wells inoculated with virus alone was plotted as the fold enhancement of infection. The mean value plus three standard deviations for three negative control wells was used as the threshold for enhancement (shown as a dotted line in A-E).

Further dilution of the immunized macaque sera to 1:100 and 1:1000 also failed to induce ADE for any of the four serotypes (**Fig. S2A+B**). The only exception to this was the sera from week 16, which appeared to cause ADE with DENV4 at both 1:100 and 1:1000 dilutions. While concerning, the absence of continued ADE at later timepoints suggested that this was not necessarily a consistent effect of the vaccine. The general ability of DENVLP-induced antibodies to avoid ADE even at this low level suggests that the circulating antibody titer in DENVLP-vaccinated individuals may remain above the threshold that can cause ADE well beyond the 1-year timeframe.

### DENVLP vaccination significantly reduces viral replication

Next, we sought to determine the protective effect of the DENVLP vaccine against a live DENV challenge. Marmosets (*Callithrix jacchus*) were selected for the live challenge studies because they develop high levels of viremia upon primary DENV infection. Marmosets were immunized either twice (in DENV2 and 3 challenges) or three times (in DENV1 and 4 challenges) with 1.25µg per serotype of all four DENVLPs or with vehicle control, then challenged five weeks after the final immunization with one of the four DENV serotypes (1 x 10^5^ plaque-forming units (PFU) each) (**Fig. 3A**). All the immunized marmosets elicited robust antibody responses against DENV as measured by ELISA (**Fig. 3B**). All control marmosets challenged with DENV1 or DENV4 developed viremia, with peak viral RNA observed at 2-4 days post-challenge via RT-qPCR (**Fig. 3C**). In contrast, all DENVLP-immunized marmosets showed significantly reduced viral RNA levels at peak infection (67,000 copies/mL for DENVLP-immunized animals vs 194,000 copies/mL for controls) (**Fig. 3C**). When total infectious virions were measured via plaque assay, control marmosets challenged with DENV1 showed peak viremia around 1.9 x 10^4^ PFU/ml at day 4, while DENVLP-immunized marmosets had only 1.2 x 10^3^ PFU/mL (**Fig. 3D**), a more than 10-fold decline in infectious virions. Similarly, the control marmosets challenged with DENV4 developed peak viremia more than 3 x 10^2^ PFU/ml, whereas VLP-immunized marmosets had no detectable plaques observed at day 4 (**Fig. 3D**). Upon challenge with DENV2 and DENV3, we failed to detect infectious virions in either immunized or control samples via plaque assay (data not shown). However, a significant increase in DENV2 and DENV3 genomic RNA copies/mL was observed in the blood of control animals via qPCR, suggesting that viral replication was occurring in unvaccinated marmosets (**Fig. 3D**). At the peak of viral RNA load in the control group, the level of DENV2 viral RNA observed in VLP-immunized marmosets was nearly 10-fold lower (2.6 x 10^4^ copies/mL for DENVLP-immunized vs 2.6 x 10^5^ copies/mL for controls) and the level of DENV3 viral RNA observed in DENVLP-immunized marmosets was more than 3-fold lower (6.8 x 10^6^ copies/mL for DENVLP-immunized vs 2.1 x 10^7^ copies/mL for controls) (**Fig. 3E**). These data suggest that immunization with tetravalent DENVLP provides significant protection against dengue infection for all four DENV strains.

**Figure 3.**
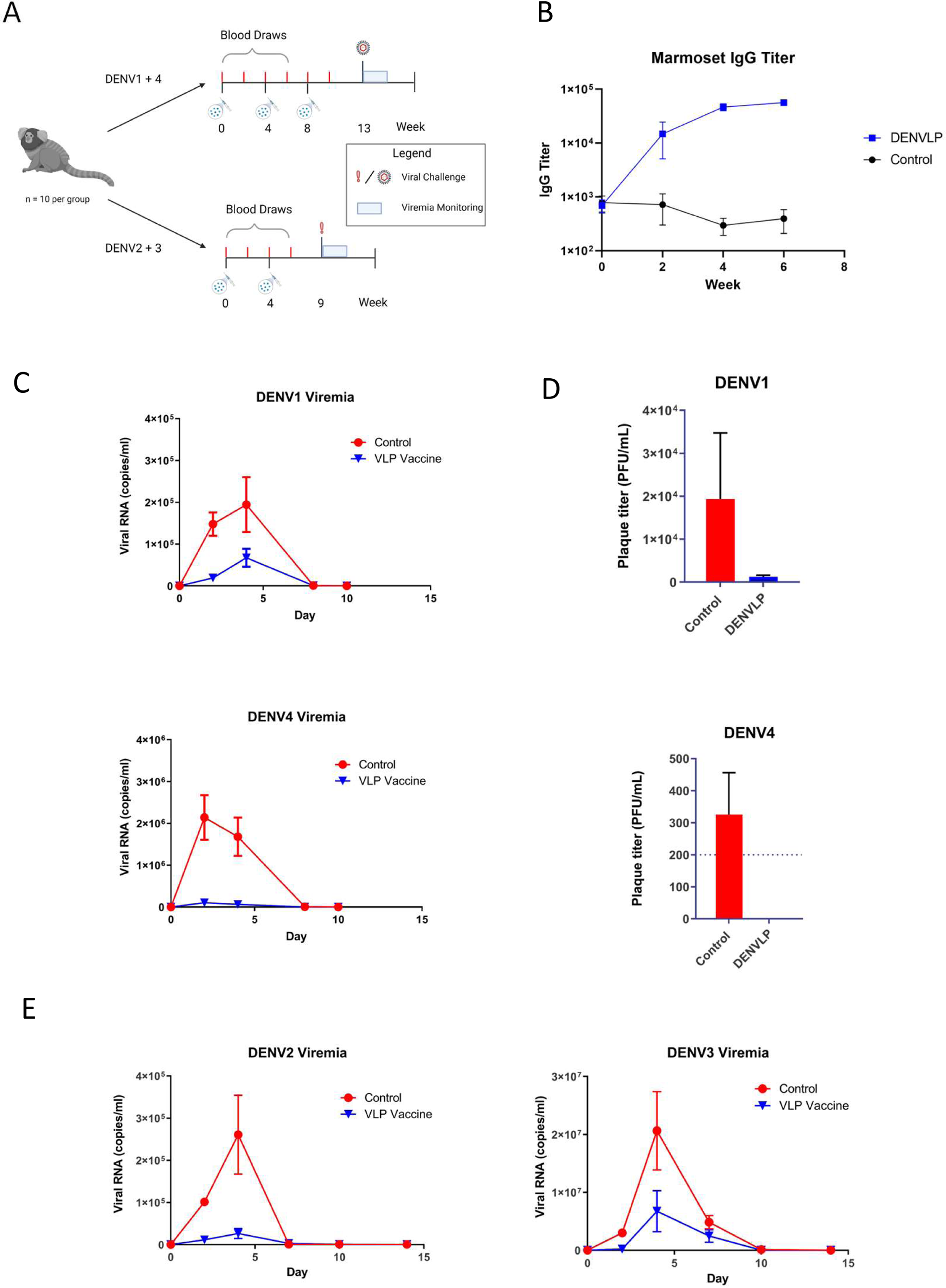
DENVLP-immunized marmosets are protected from severe viremia upon DENV1-4 challenges. **(A)** Graphic of marmoset challenge experiment. Forty marmosets were divided into four equal groups and immunized with either 5 µg tetravalent DENVLP (n = 6) or vehicle control (n = 4) either twice (weeks 0 and 4, DENV2 and 3 challenge) or three times (weeks 0, 4 and 8, DENV1 and 4 challenge) **(B)** Serum samples drawn from DENVLP- or control-immunized marmoset from weeks 0-6 were used to measure the total anti-DENV IgG titer via ELISA. **(C + E)** Measurement of blood viremia by RT-qPCR. Viral RNA was purified from marmoset blood samples drawn every two days from day 0-10 (DENV1+4) or day 0-14 (DENV2+3), and then measuring the total number of genomic copies per mL of blood using serotype-specific primers against DENV E protein. (D) Measurement of blood viremia by plaque assay. Blood samples drawn from each marmoset at day 4, the peak of mean viremia (measured by RT-qPCR), were serially diluted and incubated with Vero cells, and the mean number of plaque forming units per mL of blood was measured for each marmoset.

To determine if the protection against viral infection generated by the DENVLP vaccine was conferred primarily by the neutralizing antibodies produced, we investigated whether IgG from DENVLP-immunized macaques could protect against lethal challenge in a passive transfer animal model^23^. Purified total IgG from macaque sera drawn at either week 0 (pre-immune) or week 12 (VLP-immunized) was transferred into AG129 mice (n =4 per group), which lack interferon receptors. The mice were subsequently challenged with a lethal dose (1 x 10^6.5^ PFU) of DENV224 horus after IgG transfer Mice that received pre-immune IgG developed severe infection and died by 15 days post-infection, while mice that successfully received DENVLP-immunized IgG were protected from lethal infection, (**Fig. 4**). These results indicate that the humoral immune response induced by tetravalent DENVLP immunization confers protection against severe dengue infection.

**Figure 4.**
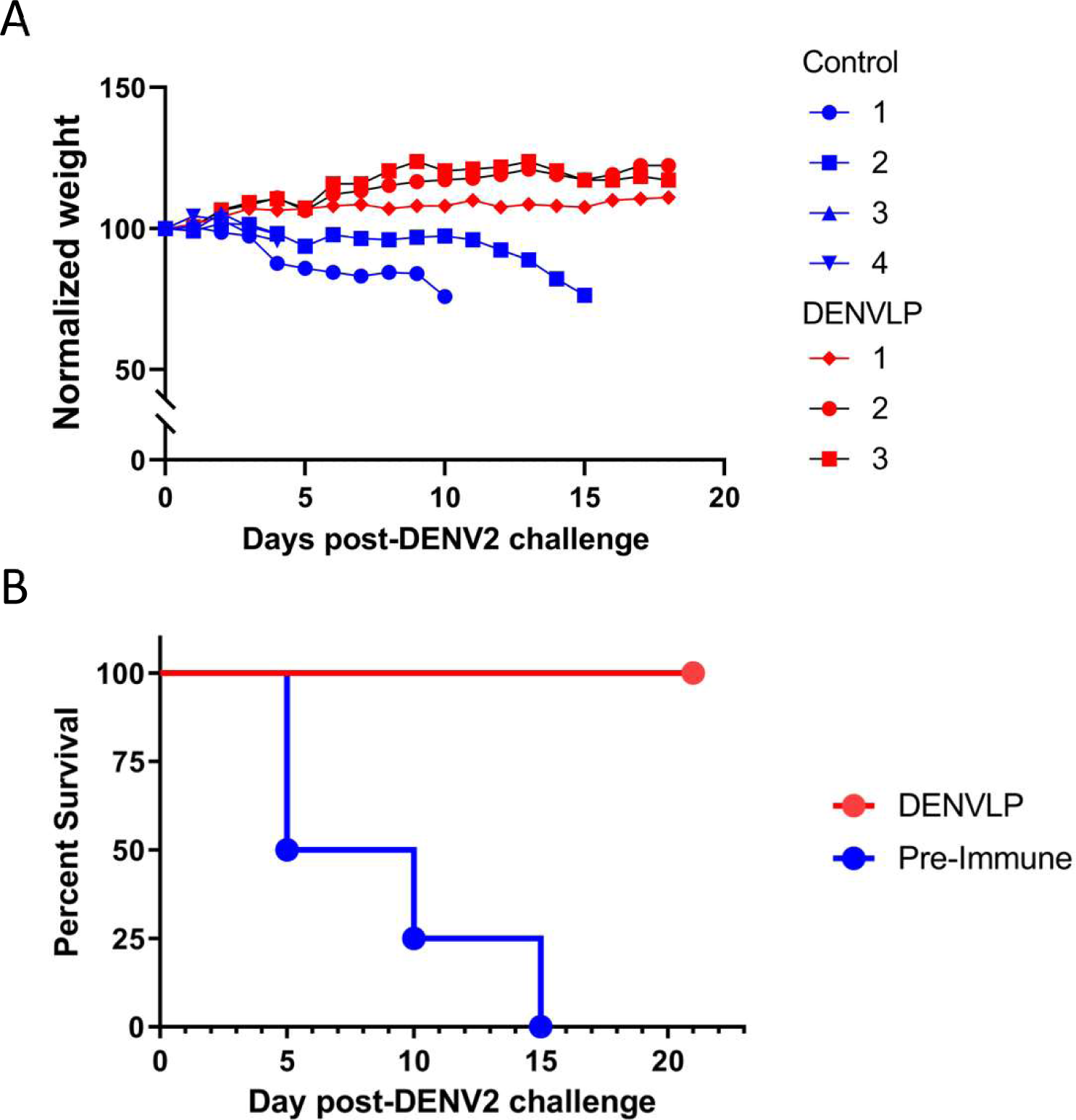
Passive transfer of DENVLP-immunized macaque sera prevents lethal DENV2 infection of AG129 mice. Total IgG was purified from blood samples drawn from all 18 macaques either prior to first immunization (pre-immune, n = 4) or at week 12 (DENVLP, n = 3), and the purified IgG was then introduced into AG129 mice via i.p injection 24h prior to challenge with a lethal dose of DENV2. (A) The weight of each individual mouse following DENV2 challenge normalized to its initial pre-challenge weight. (B) The overall survival rate of AG129 mice receiving IgG from macaques before or after DENVLP immunization.

## Discussion

To meet the growing global health challenge of dengue, a next-generation vaccine must meet several key criteria to be effective: it must be able to be used in high-risk groups, quickly generate protective immunity against all four dengue serotypes, and ensure that protection is durable over the long-term. Here, DENVLP-immunized macaques generated a robust NAb titer against all four dengue serotypes within just two weeks of immunization, with a peak NAb titer observed at four weeks after the second booster immunization, and NAb titers remained high for up to one year after initial immunization. The induction of a high NAb titer is essential to avoid a “danger zone” where anti-DENV antibodies are present at lower levels than what is required to neutralize the virions, which could leave vaccine recipients at risk of ADE until subsequent booster doses raise their NAb titers^24^. Another risk period could occur later after vaccination if the NAb titer gradually wanes into dangerous non-neutralizing levels. Critically, the anti-dengue antibodies present in macaque sera from week 2-52 failed to induce ADE for any other four dengue serotypes using a BHK-FcγR reporter system. These results suggest that the DENVLP vaccine avoided both the early and late risk periods. Both the speed and durability of the immunogenicity generated in macaques provide evidence that the DENVLP vaccine shows promise as a next-generation dengue vaccine.

All three doses of DENVLP vaccine (0.5, 5 and 50 µg total VLP) produced robust NAb titers in macaques, and at most timepoints there was no statistically significant difference between the NAb titer induced by the highest and lowest vaccine doses. These results suggest that the quantity of DENV antigen required to induce a robust immune response may be lower than even the lowest dose delivered here. For comparison, immunization of rhesus macaques with a recombinant subunit vaccine produced much lower NAb titers even when vaccinating with 250 µg total DENV E protein^25^. A plausible explanation for why the DENVLP vaccine was able to induce strong immunity with so little antigen is that the repetitive structure of VLPs allows for the robust stimulation of T- and B-cells required for a strong immune response. For this reason, VLP vaccines may be an attractive alternative to other non-live vaccination methods. However, a dose variation clinical trial will need to be conducted to establish the minimum dose of VLP vaccine required to induce durable protective immunity in humans.

It is not a perfect comparison to measure the NAb titer against two different strains of DENV generated by FRNT_50_ values, and even more difficult to compare relative NAb titers generated by different vaccines when the methods used for calculating those titers are different. However, there is some value to comparing the NAb titers for each serotype produced by tetravalent DENVLP vaccination in NHPs to the NAb titers produced by other vaccines. Prior to their use in humans, both currently approved live attenuated vaccines were first tested in non-human primates. Immunization of macaques with equal doses of each serotype of the Dengvaxia vaccine produced a robust DENV4 NAb response at 1 month after immunization, but relatively lower titers of DENV1, 2 and 3 Nabs^26^. This mirrored the overall efficacy of Dengvaxia in seronegative children in phase III studies, where the risk ratio for hospitalization with DENV4 was lower than that of DENV1, 2 or 3^27^. Similarly, immunization of macaques with a tetravalent Qdenga formulation produced an imbalanced antibody response, with a significantly higher DENV2 NAb titer than DENV1, 3 or 4 NAb titers^28^. This matched the overall efficacy of QDenga vaccination in seronegative recipients, where they showed good efficacy against DENV2 and poor efficacy against DENV3 and 4^13^. Rather than comparing the overall NAb titers measured in macaques following DENVLP immunization, it is rather better to look at the relative ratio of NAb titers. Following DENVLP immunization, there was no serotype with a consistently higher NAb titer across weeks 2-52. The NAb titers against all four DENV serotypes rose and then declined at a similar rate, indicating that the circulating antibodies were not imbalanced. The balanced NAb titers throughout the one-year period likely contributes to the absence of ADE activity that we observed. We hypothesize that both the balanced NAb titers and the absence of ADE stem from the consistent presentation of equal quantities of antigen for DENV1-4 by the VLP vaccine system. Finally, we predict that these results suggest that DENVLP vaccination carries a low risk of having a significantly different efficacy rate against different DENV serotypes.

Because of the high NAb titer generated by the DENVLP vaccine, we were able to demonstrate that this vaccine can protect against live dengue infection with all four serotypes. Marmosets are susceptible to infection by all four DENV serotypes, however vaccination with DENVLP vaccine provided significant protection against viremia, as observed by both plaque assay and qPCR. These results demonstrate that vaccination with DENVLP produced immunity capable of preventing severe infection even when challenged with high doses of live virus. The passive transfer of purified IgG from VLP-immunized macaques to immunocompromised AG129 mice provided protection against DENV2 infection, confirming that the NAbs induced by vaccination were responsible for this protection.

Dengue virus produces many immature particles during infection, and antibodies against the immature prM protein have previously been implicated as a risk factor for ADE and severe secondary infection^29^. In accordance with this risk, there may be some concern that a vaccine using the prM protein (instead of mature M protein) may increase ADE susceptibility. However, our measures of ADE activity using FcγR-expressing BHK cells demonstrate that this is not the case for any of the four DENVLP constructs. Given that the prM region of all four constructs is the same, the serotype-specific immunity must be conferred by the E proteins for each serotype. Therefore, these results suggest the neutralizing antibodies generated against the E protein are sufficient to counteract any prM antibodies that may enhance infection. Another previously reported risk factor for ADE is antibodies specific to the fusion loop region of the E protein^30^. The DENVLP constructs, which contain the F108A mutation to improve overall yield, may thereby limit ADE by generating antibodies exclusively to epitopes other than the fusion loop.

While these results are promising, there are several limiting factors that qualify our interpretation of the results. For example, while the BHK-FcγR system can indicate whether the virions taken up via Fcγ receptors have been insufficiently neutralized, the *in vitro* incubation of immunized sera and DENV means that this experiment only roughly approximates whether ADE would be prevented in human patients. Similarly, while marmosets are an effective model for DENV infection, the protection conferred by the DENVLP vaccine will need to be confirmed in a clinical trial in humans. Ultimately, these results demonstrate that a VLP-type vaccine is a promising candidate for inducing balanced, robust immunity against all four dengue serotypes, and the results support further evaluation of the safety and efficacy of these constructs in human clinical trials.

## Materials and Methods

### DENVLP preparation

The construction of DENVLP1-4 expression plasmids and production of VLPs has been reported previously^11^. Briefly, Freestyle 293F cells (Thermo Fisher Scientific) cultured in FreeStyle 293 Expression Medium (Thermo Fisher Scientific) were transfected with DENV VLP-expressing plasmids. The culture supernatant at four days after transfection was clarified, concentrated, and purified by HiTrap Q XL (GE Healthcare Life Sciences) and Foresight CHT Type II (Bio-Rad) columns with sodium phosphate gradient. Total protein concentration of purified DENV VLP was measured by Quick Start Bradford Protein Assay (Bio-Rad). Purity of the DENV VLP was assessed by SDS-PAGE followed by Coomassie dye-based staining using QC colloidal Coomassie stain (Bio-Rad). The morphologies of DENV VLPs were analyzed at the National Institute of Infectious Disease in Japan (NIID) Microscope Facility to confirm uniformity prior to their use in animal studies.

### Animal studies

All cynomolgus monkeys and common marmosets were purchased from Hamri Co. Ltd. (Koga, Ibaraki, JAPAN) and CLEA Japan, Inc. (Tokyo, Japan), respectively and housed at NIID facilities. All animal procedures were conducted in accordance with the NIID “Guides for Animal Experiments Performed at National Institute of Infectious Diseases”, approved by the Animal Welfare and Animal Care Committee of NIID, Tokyo, Japan. Sedation of animals for vaccination, infection, and blood sampling was performed with intramuscular ketamine hydrochloride (50 mg kg-1) and xylazine (3 mg kg-1).

For immunogenicity studies, 18 healthy, 4 years old male cynomolgus monkeys were used. Prior to initiation of the study, serum anti-flavi IgG titer was assessed by enzyme-linked immunosorbent assay (ELISA) to confirm that they had not previously infected with flaviviruses. VLP was mixed with Alum adjuvant immediately before intramuscular injection to femoral region at weeks 0, 4 and 8. Blood samples were taken every two weeks during the first 12 weeks, and at every four weeks from weeks 12-52. Sera samples were heat-inactivated by incubating at 60°C for 30 min. Measurement of serum anti-DENV IgG titer, neutralization antibody (NAb) titer and antibody-dependent enhancement was conducted as described below.

For the DENV viremia protection studies, forty 1 to 2-years old common marmosets (equally divided males and females) were divided into 4 challenge groups. Each challenge group was then further divided, with six marmosets receiving 5 µg total tetravalent DENVLP formulated with Alum adjuvant (DENVLP group) and four marmosets receiving vehicle solution (5% sucrose/5mM sodium phosphate, pH 7.2) as a control. The marmosets receiving DENV1 or DENV4 challenge strains received three total immunizations at weeks 0, 4 and 8, while marmosets receiving DENV2 or DENV3 challenged were immunized twice at weeks 0, and 4. For all marmosets, serum samples were taken every two weeks until DENV challenge for measurement of serum anti-DENV IgG titer. At five weeks after final immunization (week 9 for DENV2/3, week 13 for DENV1/4) all marmosets were challenged with 1 x 10^5^ PFU DENV (DENV1: 01-44 strain, DENV2: DHF0663 strain, DENV3: DSS1403 strain, DENV4: 05-04 strain) via subcutaneous injection in the back (divided between 2-3 inoculation sites). Following DENV challenge, blood was taken every two days from day 0 to day 10 (DENV1/4) or day 14 (DENV2/3) for measurement of viremia using RT-qPCR and plaque assays as described below.

### Immunogenicity assays (ELISA)

ELISA was performed as previously described^11^. Briefly, 96-well ELISA plates were coated with purified Japanese encephalitis virus (JEV, strain: JaOArS982) or DENV mixed antigens at 250 ng/well at 4°C overnight. The plates were then blocked with undiluted BlockAce (DS pharma Biomedical) for 1h at room temperature. Primary probing of the antigen was conducted using sera samples and standard control were diluted at 1:1,000. 1:1,000 diluted HRP conjugated anti-mouse IgG antibody (American Qualex) was added. Serum IgG titers were calculated from standard curve as described previously^31^. A sample titer ≥ 3,000 were interpreted as anti-flavivirus or anti-DENV IgG positive.

### Fifty Percent Focus Reduction Neutralization test (FRNT_50_)

Macaque sera from each time point were serially diluted and mixed with 40 to 50 focus-forming units of virus (DENV1 99st12A strain -genotype IV, DENV2 oost22A strain -Asian 2, DENV3-SLMC50 strain -genotype I, DENV4-SLMC318 strain-genotype I and JEV S-982 strain-genotype III). Serum-virus mixture was inoculated to Vero cell monolayer, then 1.25% methylcellulose 4000 in 2% FCS MEM was added to wells and incubated at 37°C for 3 days for DENV and 36 h for JEV. The plates were washed with phosphate-buffered saline (PBS) and then fixed with 4% paraformaldehyde solution. Then the cells were permeabilized with 1% NP-40 solution. The plates were blocked with BlockAce for 30 min and treated with 1:1,500 diluted pooled human sera having high anti-flavivirus IgG titers (32) for 1 h at 37°C. Subsequently, HRP-conjugated goat anti-human IgG (American Qualex, 1:1,500) was added and incubated at 37°C for 1 h. 0.5 mg/ml of 3,3’-diaminobenzidine tetrahydrochloride (Wako) with 0.03% of H_2_O_2_ solution was added and incubated for 10 min for staining. After washing and air drying, the number of foci per well were counted using a biological microscope. The reciprocal of the endpoint serum dilution that provided 50% or greater reduction in the mean number of foci relative to the control wells that contained no serum was considered to be the FRNT_50_ titer.

### Antibody-dependent dengue virus infection enhancement assay

FcγR-expressing BHK cell lines were seeded in to a 96 well plate and cultured in EMEM supplemented with 10 % heat-inactivated FBS and G418. Vaccine immunized macaque serum or mouse anti-Flavivirus E monoclonal antibody 4G2 were serially diluted from 1:10 to 1:10,000 with 10% FBS /EMEM, mixed with 30-50 focus-forming units of virus (same DENV strains as those used in FRNT assay) and incubated at 37°C for 1 h. Virus-immune complex were added to each well of the cells, and cultured for 48 h. The cells were washed with PBS once, dried by air, and fixed with ice-cold methanol-acetone (1:1). After the plates are air-dried, the wells were blocked with 1% horse serum in PBS for 10 min and washed with PBS for three times. The plate was then treated with anti-flavivirus antibody (mAb 4G2 at a dilution of 1:1000) at 37°C for 30 min, biotin conjugated anti-mouse IgG antibody (Vector Laboratories, 1:500) at room temperature for 30 min, and ABC (Vector Laboratories) solution at room temperature for 30 min. The plates were washed with PBS three times after each incubation step. VIP solution (Vector Laboratories) was added and incubated at room temperature for few minutes until color was developed. The plate was washed with PBS once, infected cells were quantitated (Keyence BZ-X710 microscope). Fold-enhancement values were calculated using the following formula: (mean infected cells count using FcγR-expressing BHK cells with the addition of mouse serum sample)/ (mean plaque count using FcγR-expressing BHK cells in the absence of test sample). Infection enhancement (measured as ADE activity) was tested using serum samples that was diluted from 1:10 to 1:10,000. The fold enhancement values were determined as follows: (the mean number of DENV-infected cells in wells treated with serum samples)/ (the mean number of plaques in wells with monolayers incubated in the absence of test samples). The mean value of at least 3 negative control wells plus three times the standard deviation (SD) value was used as the cut-off value to determine which samples had ADE activity.

### Measurement of viremia by RT-qPCR and Plaque Assay

The level of genomic RNA and the number of infectious virions were determined by RT-qPCR and Plaque Assays as previously described^32^. Briefly, marmoset serum was centrifuged at 200 x g to remove cells and debris, and viral RNA was extracted (High Pure Viral RNA kit, Roche Diagnostics). 5 µL of viral RNA from each sample was applied to TaqMan RT-PCR using primers and probes specific to the E protein of each DENV serotype^33^, and the resulting Cq values were used to calculate the mean viral genomic copies/mL for each marmoset at each timepoint. After calculating the day post-challenge with the highest peak viremia as measured by RT-qPCR, serum samples from the corresponding day were serially diluted 10-fold (from 1:10 to 1:10^6^) in MEM (Sigma-Aldrich) supplemented with 10% FBS. 50 µL of each dilution was added to Vero cell monolayers and incubated for 1h at 37C before removal of infectious media and replacement with MEM supplemented with 2% FBS and 1.25% methylcellulose 3000. After 72 hours incubation, cells were washed with PBS, fixed with 4% paraformaldehyde solution and then stained with crystal violet, and the total number of plaque forming units/mL of sera was calculated for each sample.

### Passive transfer experiment

A total of 15 mL sera drawn at week 0 (pre-immune) or week 12 (4 weeks after final VLP immunization) from the six macaques immunized with the highest dose of DENVLP were pooled together and purified using Melon Gel IgG Spin Purification Kit and NAb Spin column (ThermoFisher #45206, #89961) followed by buffer exchange to PBS via Slide-A-Lyzer dialysis cassettes (ThermoFisher #66012) and concentration to 25 mg/mL via Amicon ultra centrifugal filters (Sigma Aldrich #UFC9100). Purified IgG was sterilized by 0.2 µm filtration before transfer. Eight IFNα/β/γR^−/−^ (AG129) mice aged 4-5 weeks were divided equally into two groups for IP injection of 2.5 mg total IgG from either the pre-immune or VLP-immunized macaques. Mice were then challenged with 1×10^6.5^ PFU of DENV2 (strain D2S10) delivered via IP injection at 24h after passive IgG transfer. All 8 mice were then weighed daily until death, extending up to 21 days post-challenge for control animals.

## Supplementary Materials

**Figure S1.**
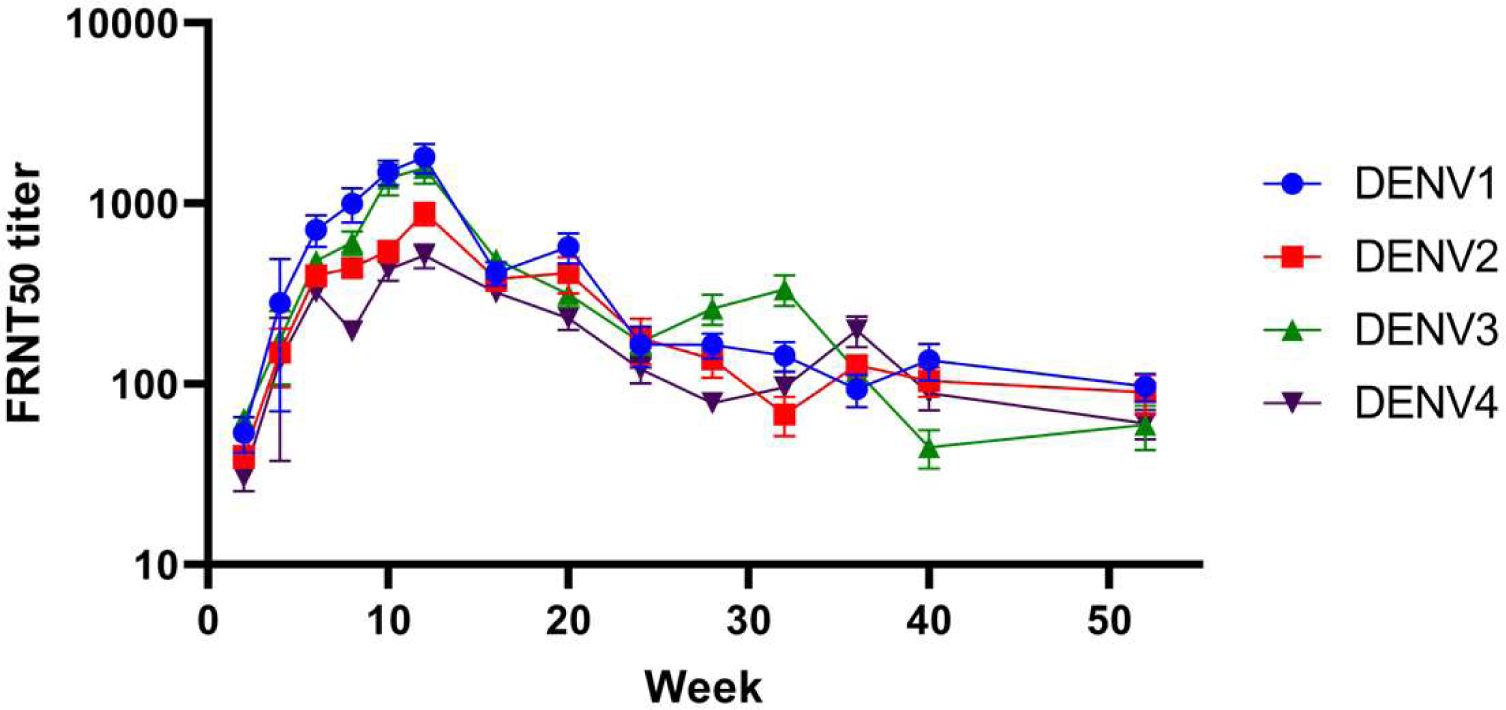
The mean neutralizing antibody titer of all 18 macaques against all four DENV serotypes. The mean neutralizing antibody titer for each serotype and time point as determined by FRNT_50_ assay. Serum samples from each macaque and timepoint were serially diluted and incubated with DENV serotypes 1-4 prior to inoculation of Vero cells. The endpoint dilution capable of neutralizing half of the viral foci relative to Vero cells incubated with DENV1-4 alone was recorded for each macaque and timepoint was recorded, and the mean value for each serotype and sample timepoint was plotted.

**Figure S2.**
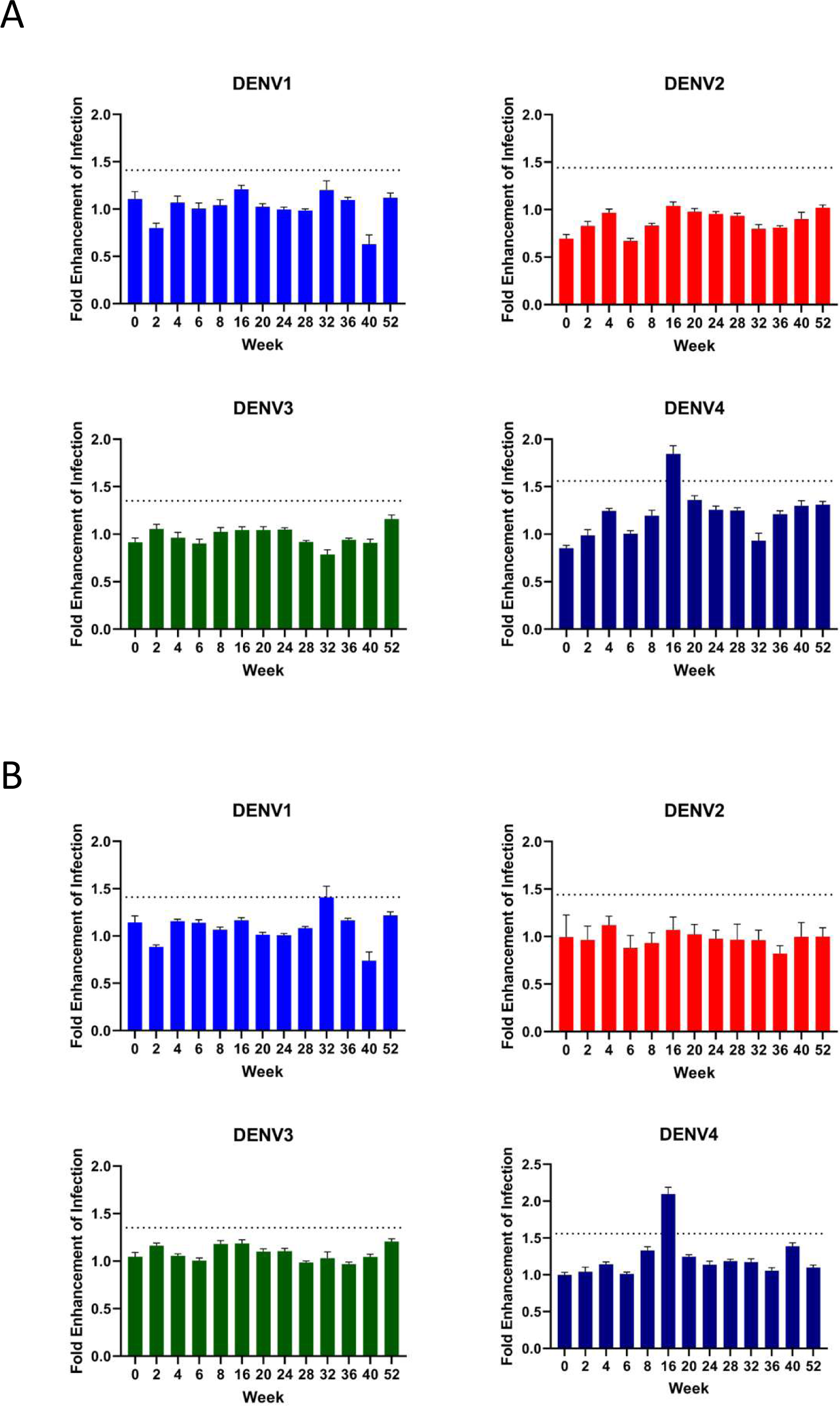
The ADE activity of macaque sera at some time points passes enhancement threshold upon further dilutions. **(A+B)** DENV1-4 were incubated with macaque sera diluted 1:100 **(A)** or 1:1000 **(B)** prior to inoculation of FCγR-BHK cells. After two days, infected cells were counted, and the ratio of infected cells in serum- or antibody-incubated wells to infected cells in wells inoculated with virus alone was plotted as the fold enhancement of infection. The mean value plus three standard deviations for three negative control wells was used as the threshold for enhancement (shown as a dotted line in A+B)

## Acknowledgements

The authors thank H. Anderson, S. Kar (Bioqual) for managing animal experiments. Figure 1B and 3A were created with BioRender. This research was funded by Global Health Innovative Technology Fund grant (G2016-109) and VLP Therapeutics. The authors declare that an intellectual property application has been filed by VLP Therapeutics based on data presented in this paper.

## Author contribution

A.U, M.M.N.T., T.N, M.L.M, Y.W, T.T.T, H.Y, Y.S, Y.A and W.A. performed the research; D.T, K.Ma, A.U, M.M.N.T., M.L.M., Y.W, M.I., H.Y, J.F.S, T.M., K.Mo., and W.A. analyzed data; D.T, K.Ma, M.M.N.T., T.N, M.L.M., Y.W. M.I, J.F.S, T.M., K.Mo. and W.A. wrote the paper. T.M, K.Mo, W.A conceived, designed, and coordinated the study. D.T, K.Ma, A.U, Y.W, and M.I are employees of VLP Therapeutics, Inc.; J.F.S is an employee and holds stocks in VLP Therapeutics, Inc. W.A is a board member, an employee and holds stocks in VLP Therapeutics, Inc. W.A is an inventor on related vaccine patent. The remaining authors declare no competing interest. All authors read and approved the manuscript.

